# Sex-specific competitive social feedback amplifies the role of early life contingency in male mice

**DOI:** 10.1101/2024.04.19.590322

**Authors:** Matthew N Zipple, Daniel Chang Kuo, Xinmiao Meng, Tess M Reichard, Kwynn Guess, Caleb C Vogt, Andrew H Moeller, Michael J Sheehan

## Abstract

Contingency (or ‘luck’) in early life plays an important role in shaping individuals’ development. When individuals live within larger societies, social experiences may cause the importance of early contingencies to be magnified or dampened. Here we test the hypothesis that competition magnifies the importance of early contingency in a sex-specific manner by comparing the developmental trajectories of genetically identical, free-living mice who either experienced high levels of territorial competition (males) or did not (females). We show that male territoriality results in a competitive feedback loop that magnifies the importance of early contingency and pushes individuals onto divergent, self-reinforcing life trajectories, while the same process appears absent in females. Our results indicate that the strength of sexual selection may be self-limiting, as within-sex competition increases the importance of early life contingency, thereby reducing the ability of selection to lead to evolution. They also demonstrate the potential for contingency to lead to dramatic differences in life outcomes, even in the absence of any underlying differences in ability (‘merit’).

## Main

Contingency (colloquially called ‘luck’ or ‘chance’) has long been recognized as an important determinant of outcomes in ecology and evolution (and to varying degrees in other fields, including philosophy, sociology, and economics (*1–15*)). The contingency hypothesis posits that an individual’s behavior, health, social position, or fitness are strongly dependent on unpredictable, uncontrollable events and experiences that occur across its life, and even in the lives of relatives and other social contacts (*5*, *16–20*). Contingent outcomes in early life are often especially important, as they can set individuals onto divergent, self-reinforcing trajectories (*1*, *2*, *5*, *17*, *18*, *20*). Recent evolutionary theory has argued that luck in an individual’s life largely swamps the importance of individual quality in determining lifetime reproductive success, and that luck in early life is especially important for such outcomes (*1*, *2*). This heightened importance of contingency in early life is consistent with the theoretical and observed limitations of phenotypic plasticity: although some plasticity is maintained throughout life, plasticity is greatest during development, and developmental decisions can restrict individuals’ future phenotypic options (*18*, *21–23*).

Many animals naturally live within larger social groups, such that contingency in outcomes is inextricably tied to individuals’ relationship to the behavior of others within societies (*5*, *24–28*). Through repeated social interactions, individuals adopt a consistent set of social phenotypes (i.e., their ‘social niche’ (*25*, *27*, *29*)). We hypothesize that competitive social processes magnify the importance of contingency in early life. For example, animals that begin with zero or small differences in competitive ability may differ in their access to resources due to variation in contingent dominance or territorial interactions (*19*, *30–33*). The resulting increased resource access for a subset of the population then improves those animals’ condition relative to those with reduced resource access, further entrenching the initial differences and magnifying the importance of early contingency (*19*, *24*, *33–36*). This process is analogous to the ‘Matthew effect’ in the social sciences, a phenomenon by which individuals or institutions that achieve early success tend to achieve ever greater success in the future (*37–39*).

Experimentally studying of the role of contingency in individual outcomes is achievable with the use of ‘replicate individuals’ that allow researchers to effectively ‘replay the tape of life’ for a single genotype under different circumstances (*40*, *41*). Studies of genetically identical animals living in the lab have demonstrated the feasibility of this approach (e.g., inbred mice: (*42–44*), inbred fruit flies: (*45*, *46*), naturally clonal Amazon mollies: (*47*, *48*)). In these studies, small between-individual differences in early behavior increase in magnitude over time, despite animals’ sharing nearly identical genetics and macro-environments (*42–48*). Yet, assessing the ways in which competitive social processes magnify or dampen the impact of early contingency requires the study of replicate individuals living under realistic, complex, dynamic social conditions— requirements that cannot be readily met under standardized laboratory conditions (*49–51*).

Here we overcome this limitation by studying the development of spatial and social behavior in age-matched, genetically identical mice living outside in a large, shared macro-environment. From infancy through adulthood, we tracked the development of the magnitude of individual differences in ecologically relevant social and spatial behaviors among genetically identical mice living under semi-natural field conditions (Fig S1).

We make two contributions to understanding the development of individuality and the role that contingency and competitive processes play in that development. First, we present the most detailed data to date on the development of semi-natural spatial and social behaviors in the prototypical biomedical model mammal. We use this data to assess the developmental timing of the emergence of individuality in genetically identical animals across a wide range of spatial and social behaviors (Fig S2). Second, we show that small initial differences in males’ ability to acquire and defend territories causes males—but not females—to enter onto divergent, self-reinforcing behavioral trajectories, with downstream impacts on a wide range of male social and spatial behaviors (Figs. 1-2). These divergent trajectories cause individual males to assume their individually distinct adult behavioral phenotypes at an earlier age than do females. Using quantitative agent-based simulations we also show that these differences between males and females can be wholly explained by sex-specific social feedback loops, which magnify the importance of early contingent experiences and shape the timing of the development of individuality (Fig 1).

**Figure 1.**
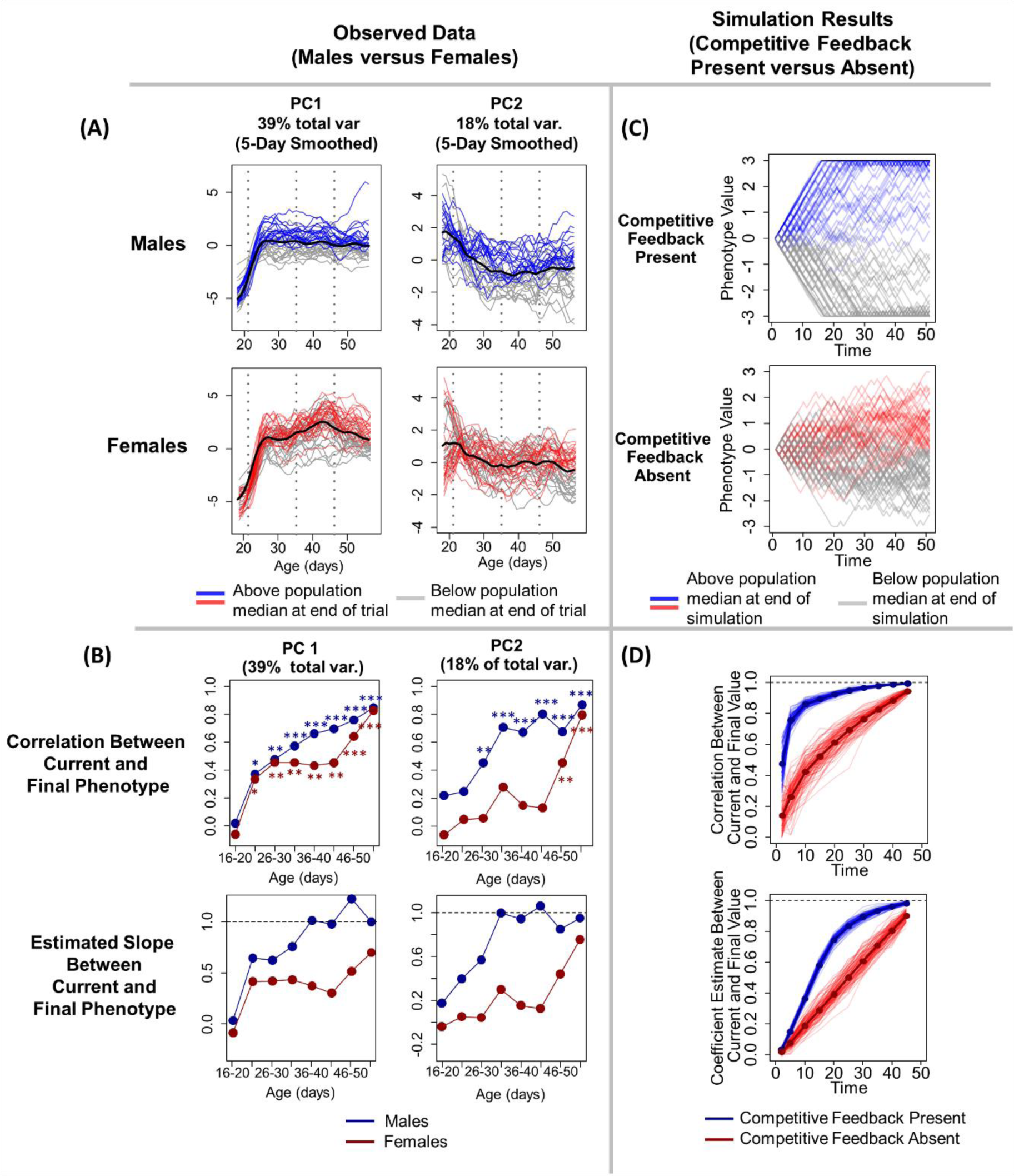
Males adopt their adult phenotypes earlier than females. (A) Traces of observed individual behavioral PC1 and PC2 values, smoothed over five days, across animals’ development. Lines are color-coded to indicate individuals’ behavior during the last three days of the experiment (age 56-58), with lines representing animals that displayed higher than median phenotype during this period in red or blue and animals that displayed lower than median phenotype in grey. (B, first row) The correlation between earlier and adult behavior is stronger in males for both PC1 (left column) and PC2 (right column). The y-axis represents the correlation coefficient between individuals’ behavior at the age-window on the x-axis and their behavior at the end of the experiment (age 56-68 days). Asterisks denote significance of the correlations depicted in each point (* p < 0.05, ** p < 0.01, *** p < 0.001). (B, second row) The slope of the relationship between earlier and adult behavior (y-axis). The slope of this relationship is consistently closer to 1 for males than for females. (C-D) Each of the observed results in (A) and (B) are closely mirrored by agent-based simulations in which simulated individuals’ phenotypes develop either in the presence (blue) or absence (red) of competitive feedback mechanisms. (C) Traces of individual phenotypes from a single run of the simulation, with shading of traces matching (A). (D) Results from 1,000 iterations of the simulation. Comparable to (B), relationships between current and future behavior are stronger when competitive feedback is present. Detailed description of the simulation can be found in the main text and methods.

**Figure 2.**
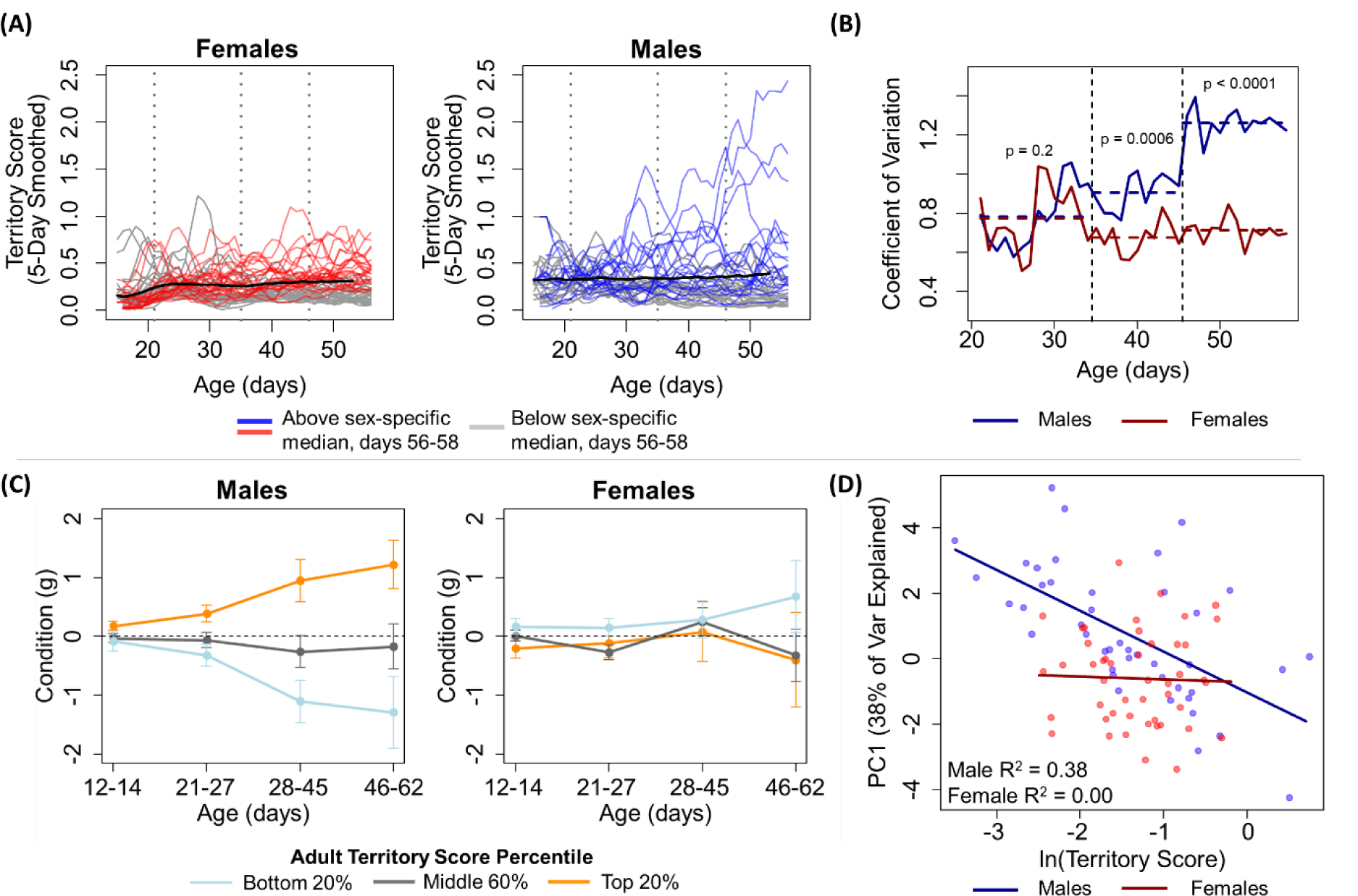
Territorial competition shapes males’ phenotypes through a sex-specific competitive feedback loop. (A) Traces of individual territory score values, smoothed over five days, across animals’ development. Lines are color-coded to indicate individuals’ scores during the last three days of the experiment (age 56-58), with lines of animals that display higher than median scores during this period in red or blue and animals that display lower than median phenotype in grey. Black lines indicate sex-specific means. Vertical dashed lines represent indicate approximate ages of weaning (‘juvenility’, 21 days), sexual maturation (‘adolescence’, 35 days), and onset of conceptive mating (‘adulthood’, 46 days) (B) Male territory scores are more variable across individuals than female territory scores, a significant difference that emerges concurrent with sexual maturity and further increases following the onset of successful mating. Horizontal dashed line segments indicate the average coefficient of variation across the juvenile, adolescent, and adult stages. (C) Male adult territory scores are predicted by small differences in body mass in early life, a difference that is magnified over time. The relationship is absent in females. The y-axes represent deviations from age-predicted body mass. (D) Adult territory score strongly predicts an integrative measure (PC1) of the 16 other spatial and social phenotypes in males but not in females.

### Males Develop Individual Adult Behavioral Phenotypes Earlier than Females

We first assessed the developmental timing of individually distinct behavioral phenotypes and whether this timing differed in males and females. In free-living C57BL/6J lab mice, males compete for territorial control and resource access while females do not (*52*, *53*). We hypothesized that this difference in competitive experiences should cause males to diverge onto self-reinforcing developmental trajectories as some males won competitive interactions and others lost. We expect this same dynamic to be absent in females.

Over seven weeks from May to June 2022, we allowed 16 litters (n = 104 pups, 90 survived to adulthood) of the C57BL/6J lab mouse strain to develop from infancy through adulthood outside in a large (∼560 m^2^) enclosure that emulates the natural foraging and social environment of commensal house mice (Figure S1A-C). We placed litters of 2-week-old RFID microchipped pups and their mothers inside nest boxes within one of 16 “resource zones” monitored with RFID antennae. We inferred periods of social overlap using an established workflow to translate RFID positional data into estimates of the duration of social aggregations within each of the 16 monitored resource zones (Figure S1D, see (*52*) for details). Based on a total of 7.4 million RFID reads, we traced the development of 17 social and spatial phenotypes from infancy through adulthood (∼6.5 weeks, 14-58 days of age, Table S1), after which we trapped out the mice and terminated the experiment.

We first leveraged our detailed behavioral dataset to answer an outstanding question in social behavioral ecology—whether genetically identical animals display distinct, individually repeatable social behaviors under ecologically relevant contexts and, if so, when these differences emerge (*40*, *41*, *47*, *48*, *54*, *55*). We measured repeatability as the proportion of a phenotype’s total variation in each sex that was explained by individual identity (*56*) over a sliding five-day age window, after controlling for maternal/litter identity. In total we assessed 17 phenotypes including basic measures of movement patterns (e.g., the number of nightly resource zones that an animal visited), measures of social phenotypes (e.g., the nightly number of opposite-sex animals encountered and territory scores), and derived social network measures (e.g., eigen vector centrality, see Table S1 for a description of phenotypes). We detect significant repeatability in all measured phenotypes, for both sexes, with repeatability emerging as early as age 21 days (Fig S2), roughly 1 month earlier than reported for spatial behavior of female populations of this strain in enriched lab *vivaria* (*11*, *30*). Animals’ behavior was repeatable prior to sexual maturity (∼age 35 days) for 15 of 17 phenotypes in males and 16 of 17 phenotypes in females (range = 21-55 days, median = 26 days, see Fig S2 G-H).

Having established that genetically identical mice still displayed strongly repeatable individual suites of behavior, we next assessed the developmental timing at which males and females assumed their individually distinct adult behavioral phenotypes. To generate an integrative measure of animals’ spatial and social behavior, we used principal component analysis to reduce the dimensionality of 16 of our 17 behavioral phenotypes into two principal components that accounted for a majority of the total variation in our dataset (57% total across PC1 and PC2; ‘time of first nightly transition’ phenotype could not be included because values were missing before animals began moving between zones). Many phenotypes were loaded onto PCs 1 and 2 without any one phenotype being particularly influential (see Table S2 for full loading information).

We identified animals’ final adult behavioral phenotypes by taking the average of each individuals’ PC1 and PC2 scores during the last three days of the experiment (age 56-58 days, Fig 1A). For each sex we then assessed the relationship between individuals’ phenotypes at earlier time points and these final adult behavioral phenotypes by building linear regressions between final adult phenotype and individuals’ average phenotypes over five-day, non-overlapping windows (e.g. age 21-25 days, 26-30 days, etc.). From each of these models we extracted the correlation between earlier and adult phenotype and the estimated slope of the relationship between earlier and later phenotypes (Fig 1B).

For both PC1 and PC2, males assumed their individually distinct adult behavioral phenotypes earlier than females. Male behavior became predictive (p < 0.05) of final adult behavior at or before females (PC1: 26 days vs. 26 days, PC2: 31 days vs 46 days, Fig 1B). The strength of the correlation between earlier and later behavior for the same individual was also substantially higher for males than for females across most or all of development (PC1: days 31-50; PC2: days 16-50, Fig 1B). Males’ behavior at earlier ages also more closely aligned with their adult behavior in absolute terms. That is, the slope of the linear relationship between earlier behavior and adult behavior was closer to 1 for males than for females, and this strong relationship developed at an earlier age for both PC1 and PC2 (Fig 1B). We performed this principal component analysis using all daily phenotype data for both males and females, but all of the above results hold if we instead generate separate sex-specific PC values (Figure S3)

To aid in interpretation of Figure 1B, we provide an example of one set of models (Figure S4), comparing the strength of the relationship between behavior immediately after sexual maturation (ages 36-40 days) and individuals future behavior in adulthood (days 55-58) for males and females. Here the strength of the relationship is much stronger for males for both PC1 (male R^2^ = 0.43; female R^2^ = 0.18) and PC2 (male R^2^ = 0.44; female R^2^ = 0.02). And the slopes of the relationship between earlier and later behavior is much closer to 1.00 for males than for females for both PC1 (male estimate = 1.01, 95% CI = 0.6-1.4; female estimate = 0.37, 95% CI = 0.1-0.6, non-overlapping with males) and for PC2 (male estimate = 0.95, 95% CI = 0.6-1.3; female estimate = 0.2, 95% CI = −0.2-0.5, non-overlapping with males).

We next assessed whether these sex differences in the developmental timing of individuality could be fully explained by sex-specific difference in the importance of competitive feedback in amplifying the importance of early contingency. To do so, we built a quantitative agent-based model to generate empirical predictions of how competitive processes shape the long-term phenotypic impacts of early contingency (Fig 1C-D). In this model we assumed that all individuals start with the same value of a phenotype. Individuals’ phenotypic value then changes at discrete timesteps across their lives, with the nature of that change depending on the presence or absence of competitive feedback. In simulations where competitive feedback was absent, the direction of change in phenotypic value was randomly chosen to be positive or negative. When competitive feedback was present, an individual’s phenotype increased in value if it won a competitive interaction with a randomly chosen individual and declined if they lost. The probability of winning the interaction was dependent on the relative phenotypic values of the two interactants (see methods for complete details).

The results of the simulation closely mirror our observations of differences in the development of individuality in males and females in our system. Specifically, when competitive feedback loops are present (i) the correlation between behavior at any given time and behavior at the end of the modeled period is stronger, and (ii) the slope of the relationship between earlier and later behavior is closer to 1.0. Thus, it appears that the sex-difference that we observe in the development of behavioral individuality could be entirely explained by differences in sex-specific competitive processes that amplify contingent early life differences in phenotype.

### Territoriality acts as a sex-specific competitive feedback loop

We next assessed whether males and females displayed differences in the strength of resource competition in a fashion that would support this putative sex-biased competitive feedback loop indicated by the model analysis in Figure 1. To do so, we estimated individuals’ nightly resource access by calculating a nightly territory score for each animal (see methods). Consistent with males and females experiencing different levels of competition for resource access, territory score varied more among males than it did among females (Fig 2A). This difference emerged concurrently with the onset of sexual maturity, the period when we expect intrasexual competition to increase in intensity. Although variation in territory score is comparable for males and females during the juvenile period (ratio of variance = 1.5, p = 0.2, two-sided F test), following the onset of sexual maturation (∼age 35 days), males displayed significantly higher variation than did females (ratio = 2.9, p = 0.0006), a difference that further increased following the onset of successful mating (∼age 46 days, ratio = 3.6, p < 0.0001, Fig2A).

Two additional pieces of evidence are consistent with males, but not females, experiencing strong competitive feedback that set them on self-reinforcing divergent life trajectories. First, small individual differences in early body mass (measured, days 21-27) predicted adult (days 46-58) territory scores for males (p < 0.05, see Fig 2C, Fig S5) but not for females. Prior to release into the enclosure, very minor differences in infant body condition did not predict future territory access in either sex (days 12-14 compared to adult, p > 0.05, Fig S5), consistent with individuals starting out on an approximately ‘even playing field’. The magnitude of the differences in body mass between males with differential resource access then increased over development (age x adult territory score interaction: p = 0.004, Fig 3C, S5), consistent with males experiencing a competitive feedback loop that increased the condition of winners relative to losers. This developmental pattern was absent in females (p = 0.19, Fig 3C). In adulthood, this mass measure in females is partially confounded by pregnancy status, but at no point (even before any pregnancies began) was there a relationship between body condition in females and adult territory status (unlike in males, Fig 2C).

Second, male territory scores strongly predicted the rest of males’ behavioral phenotypes, while the same was not true of females, indicating that our measure of competition in males has major impacts on males’ daily behaviors and ability to reproduce. Specifically, we performed a principal component analysis using the other 16 phenotypes that we measured in adulthood (i.e., excluding territory score) to obtain a single integrative measure of animals’ other behavior (PC1 explained 41% of the total variation in this dataset). Males’ adult territory scores strongly predicted this adult PC1 value (Fig 2D, p < 0.0001, R^2^ = 0.38), while females’ territory scores did not (Fig 2D, p = 0.86, R^2^ = 0.00). The same conclusion holds if we assess individual adult phenotypes, rather than this principal component measure (see Table S3 for all 16 comparisons). Individuals’ access to members of the opposite sex provides a particularly striking, fitness-relevant example from this more granular analysis: for males, the average number of females met on a given night is very strongly, positively associated with territory score (R^2^ = 0.67, p < 0.0001), while territory score does not predict the number of males met by females (R^2^ = 0, p = 0.7).

Thus, male territorial competition appears to act as a sex-specific competitive feedback loop. Small initial differences in male body mass became magnified over time, depending on territorial control. Male territorial control then had downstream implications for a wide range of ecologically and fitness relevant spatial and social behaviors.

## Discussion

Our results provide empirical support for the hypothesis that contingency (or ‘luck’) in early life can have a major and sex-specific impact on the development of animals’ individual differences in social and fitness-relevant phenotypes. Sex-specific competitive feedback loops magnify the importance of contingency experienced early in life, such that young free-living male lab mice enter onto divergent, self-reinforcing developmental trajectories. Our interpretation of our empirical results is supported by their match to an agent-based simulation of expected differences in the developmental timing of individuality in the presence and absence of competitive feedback. We expect the sex-specificity of such competitive feedback loops to vary across different species, depending on a given species’ specific social behavioral ecology. For example, in hyenas and other female-dominant species, we would expect the reverse pattern to be present, such that females’ outcomes to be more dependent on early luck (e.g. the relative social status of the matriline into which they were born, (*57*, *58*)) than males’ outcomes.

Our results suggest an inevitable limitation of sexual selection to shape behaviors. Intrasexual selection relies on within-sex competition resulting in differential reproductive success, and for variation in this success to be heritable (*59–61*). But here we have shown that intrasexual competition also magnifies the importance of contingency in later life outcomes in the sex expressing that competition. As the importance of luck in determining individual phenotypic outcomes increases, selection’s ability to cause evolution declines (*1*, *2*). Thus, intrasexual selection may be self-limiting, as an increase in the importance of competition in a single sex leads in turn to an increase in the importance of contingency in determining individual outcomes. And to the extent that intersexual choice is at least partially dependent on intrasexual competition, we expect the increased importance of luck to act as a limit on the effectiveness of intersexual selection as well. This increased importance of luck in systems with intrasexual competition may help to explain why sexual selection fails to fully deplete genetic variation, despite strong selection imposed by mate choice and intrasexual competition (i.e., the lek paradox (*62–64*)).

Our results provide a strong biological analog to the Matthew Effect, an often-observed phenomenon in social science whereby small individual advantages earlier in life are correlated with ever larger advantages over time (*37–39*). Such processes are understood to be the result of social feedback mechanisms, by which an individual’s initial success improves their opportunities for future success as well as the perception by other members of society of the individual’s potential for success (*39*, *65*). The extent to which Matthew Effects are specific to human societies has remained an open question (*39*). Our results suggest that Matthew Effects (i) may have a biological origin, (ii) are especially likely to occur in highly competitive environments or among groups that face high levels of competition, and (iii) may emerge even in the absence of any variation in underlying individual quality or ability.

The sources of inequality in human society are of central interest to both moral philosophy and public policy (*66–68*). As with reproductive success in non-human animals, human outcomes are likely to be partially explained by differences in genotypes (*69*). However, we show here that even among isogenic animals, individuals still attain dramatically different phenotypic and fitness outcomes. Our results add to sociological and biological literature that underscore the potential importance of unpredictable, uncontrollable experiences in generating differences in outcomes even when differences in underlying quality (or ‘talent’) are small or non-existent (*12*, *15*, *16*).

## Acknowledgments

We gratefully acknowledge our sources of funding that made this work possible. MNZ has been supported by an NSF postdoctoral fellowship in biology (award # 2109636) and a Klarman postdoctoral research fellowship from Cornell University. CCV is supported by a Mong Neurotechnology Fellowship from Cornell University. This work was also supported by Pilot and Feasibility awards to MNZ and MJS from the Animal Models for the Social Dimensions of Health and Aging Network (project #5R24AG065172-03). The costs of care for the mouse colony were supported in part by R35 GM138284 to Andrew Moeller.

## Data and Code Availability

While this manuscript is under review, all data and code supporting this manuscript and its analyses can be found in this Box folder: https://cornell.box.com/s/9jz4hp4kl1wnxhb5u3q275m650kfub7x

## Supplementary Materials

Materials and Methods

Supplementary Text

Tables S1 to S3

Figs. S1 to S5

## Materials and Methods

### Field Enclosure and RFID Data Collection

A detailed description of the enclosures at Cornell University’s Liddell Field Station can be found elsewhere (*52*), so here we only describe those elements critical to the success of this experiment. The enclosure is 15m x 38m, approximately 9,000 times the area of a typical mouse cage. Within the enclosure we set up 16 plastic tubs (31 gallon storage totes, Rubbermaid, USA), placed into four neighborhoods of (Figure S1E). Each tub (hereafter “resource zones”) contained *ad libitum* food access along with a nestbox that provided insulation and shelter from adverse weather conditions. We equipped each zone with a single joint entrance/exit made out of a 6-inch-long PVC pipe (2” in diameter). These resources and the single entrance made the resource zones highly valuable, defendable areas that are meant to mimic the foraging landscape of commensal mice. To track the comings and goings of mouse visitors to each zone, we placed a 10-inch RFID antennae (Biomark, USA), beneath the entrance tube of each zone. The antennas were connected to a central monitoring system (Small Scale System, Biomark, USA) and transmitted RFID reads at a rate of 2-3 Hz.

### Study Subjects

We bred 16 litters of C57BL/6J mice by pairing 9-week old virgin males and females that we ordered from Jackson Laboratory (Bar Harbor, ME). We timed pregnancies such that all litters were born within 48 hours of each other, allowing infants to be approximately age-matched. Males were removed from breeding cages two weeks after pairing to prevent mothers from becoming pregnant again following parturition.

When pups were 8-10 days of age, we anesthetized litters and their mothers using isoflurane and injected either 1 (pups) or 2 (mothers) PIT tags (Biomark Mini HPT10) subcutaneously. When animals were 12-14 days old (12: n = 49 pups, 13: n = 51 pups, 14: n = 4 pups) we placed litters along with their mothers and their nesting material in cardboard transfer containers. We then transferred litters and mothers to our field enclosures and placed them inside of nest boxes within resource zones. We balanced litter sizes across the four neighborhoods, placing 26 pups and 4 mothers in each neighborhood. 3 neighborhoods had litter sizes of 5, 6, 7 and 8 pups. The final neighborhood had litter sizes of 4, 6, 7, and 9 pups.

We then allowed animals to develop and live largely undisturbed (but see “Mass Measures” below) for 46 days, at which point we terminated the experiment. We selected this length of time to prevent animals from giving birth in our enclosure. In total, 85 of the 104 pups that we placed outside survived until the last three days prior to the end of the experiment (survival rate = 82%). These animals’ data were included in all analyses. 5 other animals survived until at least 46 days of age (the onset of conceptive mating). These animals (n = 90 total) were included in all analyses except those appearing in Figure 4. An additional 5 animals (n = 95 total) survived until 30 days of age, and we included these animals’ data in the repeatability analyses presented in Figure 2.

### Mass Measures

When pups were all 21-23 days of age, we caught by hand all living animals in the enclosure to measure weaning mass. At this time, we also removed half of the mothers living within the enclosure. This manipulation was performed to assess whether our animals relied on post-weaning maternal care. Though we do not discuss these results in detail here, there was no impact of this manipulation on dispersal, survival, juvenile behavior, or adult behavior. Following mass measures, all other animals were returned to the resource zone in which they were found.

For the remainder of the experiment, we opportunistically caught animals by hand and took mass measures. For five days a week we caught all animals present within and immediately below resource zones from a single neighborhood. We rotated neighborhoods each day to try to prevent discouraging animals from using the resource zones as a result of frequent disturbance.

On two occasions we re-captured an animal who had lost its PIT tag. To prevent these animals from making unmeasured contributions to the social environment, we humanely euthanized these animals. On one occasion we re-captured an animal that appeared to be in poor physical condition. We humanely euthanized this animal to prevent future suffering. All other animals were returned to the resource zone in which we found them.

At the conclusion of the experiment, we placed 48 Sherman live traps in resource zones for three nights until all animals were successfully caught (all animals aged 61-64 days at time of trapping). In the mornings following trapping we humanely euthanized animals, took their body masses for a final time and then dissected and preserved a range of tissues for future analysis. We placed fewer traps than there were animals due to logistical constraints on the number of daily dissections that we could perform.

### Contingency Model (*Fig 1C-D*)

To assess how the presence or absence of competitive feedback magnifies or dampens the importance of early-life contingency on later life outcomes we built a probabilistic, agent-based model.

We simulated populations of 100 replicate individuals that each began with an identical value (0) of a given phenotype. We assumed that the phenotype had a range of possible values, which we took to be [-3,3]. Individuals’ phenotypes then changed contingently over a series of time steps. At each time step, phenotypic values for each individual increased or decreased by a value drawn from a normal distribution centered around 0.2 with a standard deviation of 0.02. Individuals continued to change their phenotypic value for 50 time steps, at which point the simulation ended.

At each timestep, half of the individuals in our populations increased their phenotypic value and the other half decreased their phenotypic value. Deciding the direction of these changes proceed by the following steps. First, at each timestep the 100 individuals in a population were randomly placed into pairs. Deciding the direction of phenotypic change for each member of the pair depended on the presence or absence of competitive feedback in the population.

In the absence of a role for competition in the development process, the individual whose value increased was selected at random. This approach is meant to simulate the impact of short-term contingency, with individuals increasing or decreasing their phenotype as a result of recent differences in the animal’s internal state or by recent non-social environmental experiences (*24*).

In the presence of a competitive feedback loop, the identity of the animal whose phenotype increased was determined by a contest. The probability of an animal winning the contest was dependent on the difference in phenotypic values of the two animals in the pair.

Specifically, the probability of the first animal in a pair winning a contest was:

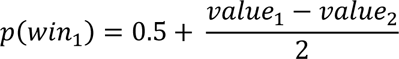

With *p(win1)* being truncated at 0 and 1. Thus, any contest between animals whose difference in phenotypic values was greater than or equal to 1 had a deterministic outcome. Contests that had smaller differences in values between contestants were probabilistic.

This approach is meant to model a competitive phenotype that determines access to resources. The probability of an individual’s phenotype increasing or decreasing depends on the outcome of a contest, which in turn depends on the value of its phenotype compared to another individual in the population. The winner of this contest acquires additional access to resources, which in turn increases its competitive ability and its phenotypic value (thus generating a competitive feedback loop (*19*, *34*)).

To generate the data in Figure 1D, we built linear models in which each individual’s final (time = 50) phenotype value was the response variable and the predictor variable was each individual’s phenotype value at an earlier point in time (times = {2, 5, 10, 15, 20, 25, 30, 35, 40, 45}). We then extracted the correlation coefficients and slope estimates from the relationship between phenotype values at these different points in time and individuals’ final phenotypes.

### Statistical Analyses

#### Analysis of Acquisition of Adult Behavior in Males and Females (*Fig 1*, S2-S3)

We first performed a principal components analysis to reduce the number of behavioral phenotypes in our population to two variables that explained a majority of the total variation in our dataset (PC1: 39% of variation explained, PC2 18% of variation explained). In this principal component analysis, we included 16 of our 17 measured daily phenotypes for all individuals from ages 14-58 days.

We then assessed the relationship between individuals’ behavior during a given period and their eventual final behavior in the experiment (measured during the last three days for which the youngest animals were present, age 56-58 days).

As in the agent-based simulation (above), we built a series of linear models for each sex where the response variable was each individual’s average PC1 or PC2 value during age 56-58 days and the predictor variable was each individual’s average value during a series of non-overlapping 5 day periods (16-20, 21-25, 26-30, 31-35, 36-40, 41-45, 46-50 and 51-55 days). We extracted the correlation coefficients and slopes of the relationships between earlier and later behavior.

#### Repeatability Analyses (*Fig 3*, S2)

We used the function rptGaussian (packaged ‘rptR’ (*70*)) to calculate repeatability estimates for behavioral measures. We calculated sex-specific repeatability across sliding 5-day age windows. We included pup age as a fixed effect and maternal/litter identity as a random effect in models with which repeatability was estimated. Thus, our repeatability estimates are the proportion of the variance in a five-day dataset of a given behavioral measure that is explained by pup identity, after controlling for pup age and maternal/litter identity.

To measure the age at which repeatability in a phenotype emerged (Figure 2C), we identified the earliest age at which animals displayed significant (p < 0.05) repeatability in a given phenotype and then continued to be repeatable thereafter for the rest of the experiment.

#### Territoriality Analysis (*Figure 2*, S5)

We calculated a nightly territory score for each animal. To do so, we calculated the proportion of sex-specific nightly RFID reads at a given resource zone that originated from each animal. We then summed these values across each of the sixteen resource zones for each animal to calculate a measure of total nightly resource access. For example, if there were 5000 male-sourced RFID reads at Resource Zone 1 on night 30 of the experiment and 4500 of them came from Male 1, Male 1 was assigned a value of 0.9 for Resource Zone 1 for that night. If Male 1 also visited exactly one other zone, where he accounted for 40% of the male-sourced RFID reads at that zone, his total value of this measure for the night (hereafter ‘territory score’) would be 1.30.

We measured coefficients of variation in territory scores for each day from age 15 to 58. To assess differences in variance in males’ and females’ territory scores we calculated each individuals’ average territory score across three different periods: (i) age 21-34 days, (ii) age 35-45 days, and (iii) age 46-58 days. We then performed a difference in variance test for square-root transformed average territory score values for the two sexes during each of these three periods (‘var.test’ function).

For each sex we then built a mixed effects linear model (function ‘glmmTMB’ (*71*)) with body condition as the response variable, predicted by the interaction between the age at which mass was measured and the log of animals’ average territory score during age 46-58 days (Fig 3C). We also included a random effect of animal ID, with age nested within ID as a random slope.

Finally, we performed a principal components analysis using all behavioral data from the 46-58 day-old period, excluding territory score. We then built linear models for each sex in which the response variable was an individual’s first principal component value (PC 1 explained 38% of the total variance in the dataset) and the predictor variable was the natural log of the individual’s average territory score in adulthood.

## Supplement

**Table S1.**
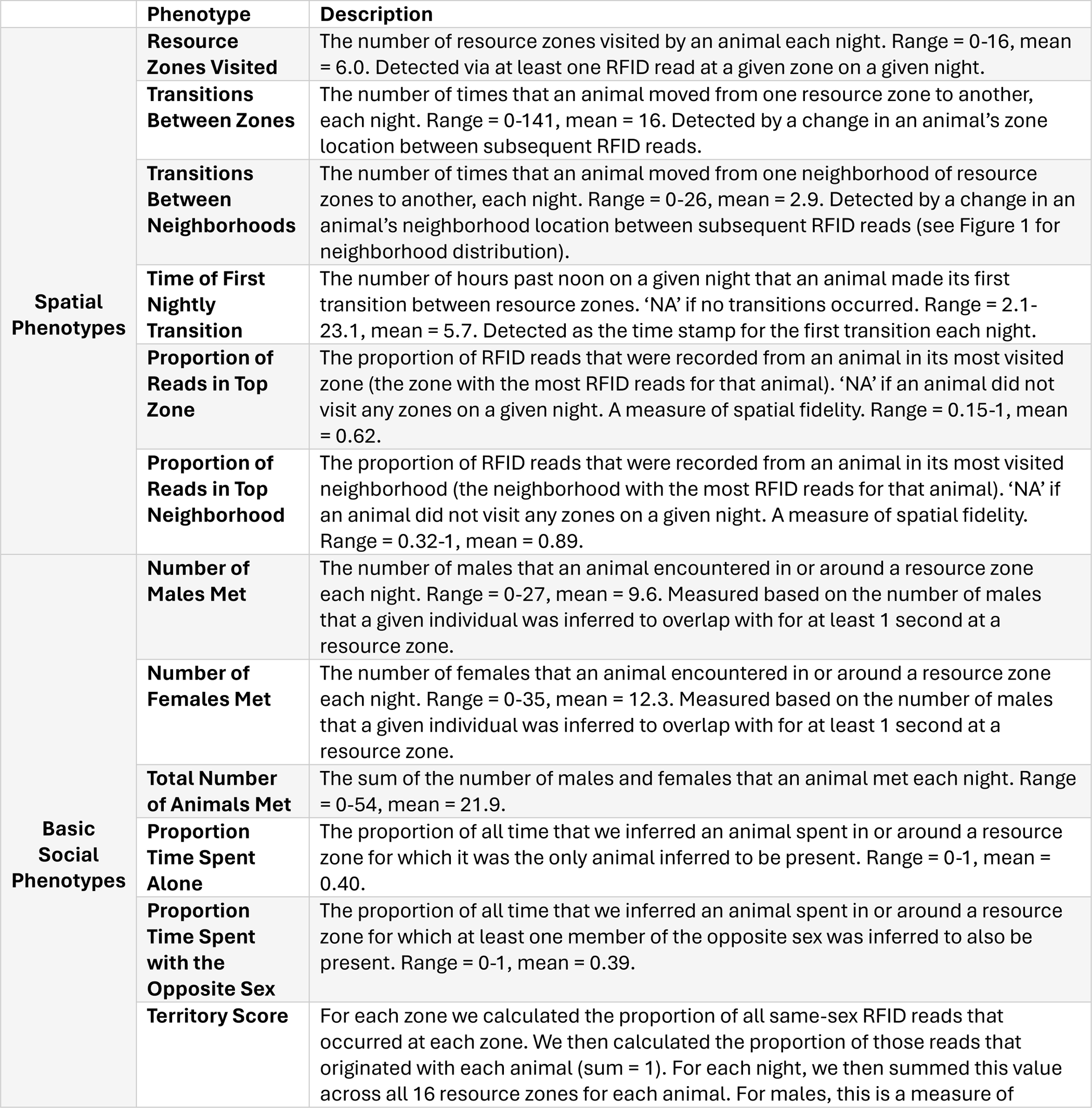
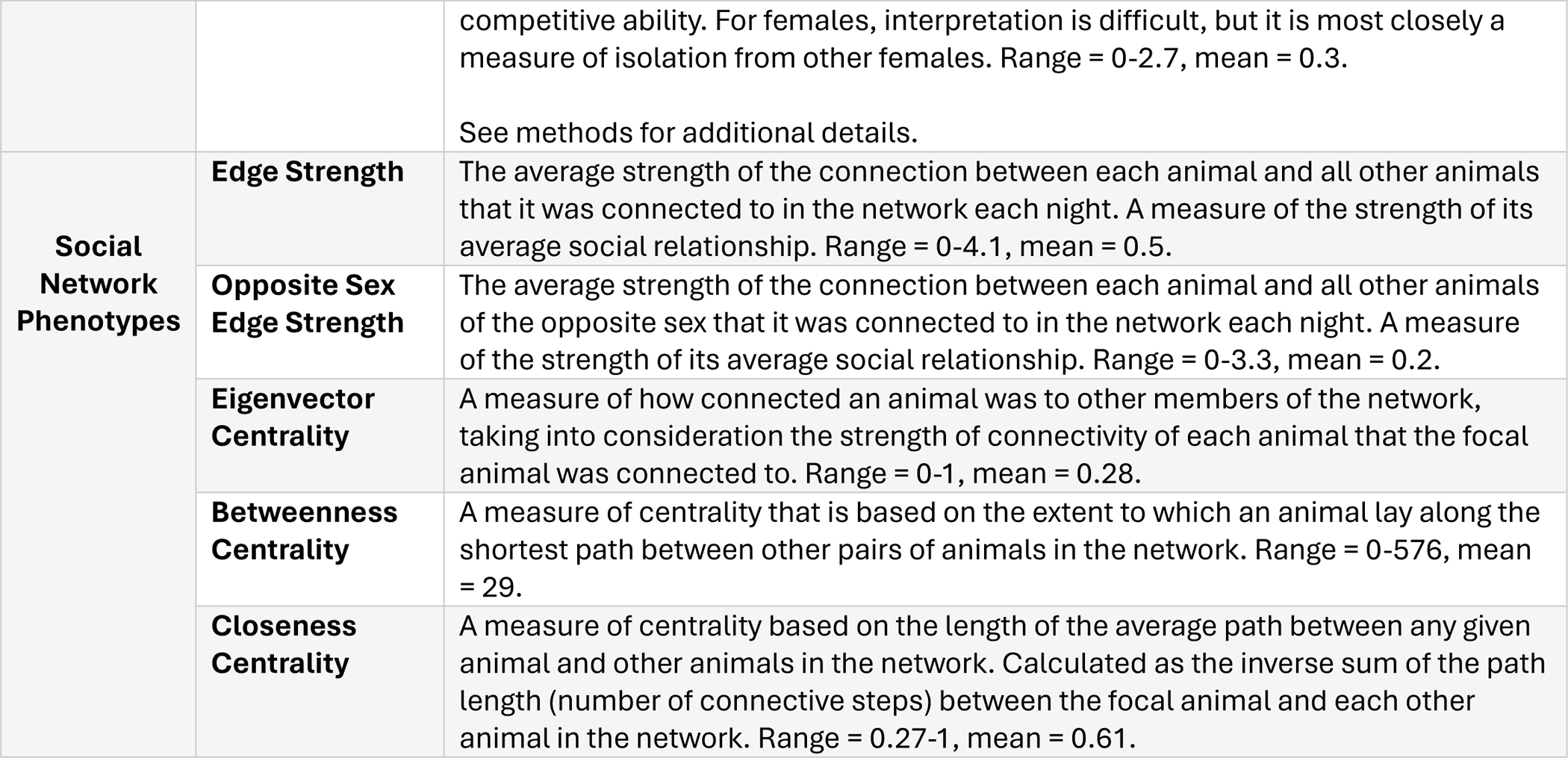
The spatial and social phenotypes measured on a daily basis in our animals, from age 15 to 58 days.

|  | Phenotype | Description |
| --- | --- | --- |
| **Spatial Phenotypes** | **Resource Zones Visited** | The number of resource zones visited by an animal each night. Range = 0-16, mean = 6.0. Detected via at least one RFID read at a given zone on a given night. |
|  | **Transitions Between Zones** | The number of times that an animal moved from one resource zone to another, each night. Range = 0-141, mean = 16. Detected by a change in an animal's zone location between subsequent RFID reads. |
|  | **Transitions Between Neighborhoods** | The number of times that an animal moved from one neighborhood of resource zones to another, each night. Range = 0-26, mean = 2.9. Detected by a change in an animal's neighborhood location between subsequent RFID reads (see Figure 1 for neighborhood distribution). |
|  | **Time of First Nightly Transition** | The number of hours past noon on a given night that an animal made its first transition between resource zones. 'NA' if no transitions occurred. Range = 2.1-23.1, mean = 5.7. Detected as the time stamp for the first transition each night. |
|  | **Proportion of Reads in Top Zone** | The proportion of RFID reads that were recorded from an animal in its most visited zone (the zone with the most RFID reads for that animal). 'NA' if an animal did not visit any zones on a given night. A measure of spatial fidelity. Range = 0.15-1, mean = 0.62. |
|  | **Proportion of Reads in Top Neighborhood** | The proportion of RFID reads that were recorded from an animal in its most visited neighborhood (the neighborhood with the most RFID reads for that animal). 'NA' if an animal did not visit any zones on a given night. A measure of spatial fidelity. Range = 0.32-1, mean = 0.89. |
| **Basic Social Phenotypes** | **Number of Males Met** | The number of males that an animal encountered in or around a resource zone each night. Range = 0-27, mean = 9.6. Measured based on the number of males that a given individual was inferred to overlap with for at least 1 second at a resource zone. |
|  | **Number of Females Met** | The number of females that an animal encountered in or around a resource zone each night. Range = 0-35, mean = 12.3. Measured based on the number of males that a given individual was inferred to overlap with for at least 1 second at a resource zone. |
|  | **Total Number of Animals Met** | The sum of the number of males and females that an animal met each night. Range = 0-54, mean = 21.9. |
|  | **Proportion Time Spent Alone** | The proportion of all time that we inferred an animal spent in or around a resource zone for which it was the only animal inferred to be present. Range = 0-1, mean = 0.40. |
|  | **Proportion Time Spent with the Opposite Sex** | The proportion of all time that we inferred an animal spent in or around a resource zone for which at least one member of the opposite sex was inferred to also be present. Range = 0-1, mean = 0.39. |
|  | **Territory Score** | For each zone we calculated the proportion of all same-sex RFID reads that occurred at each zone. We then calculated the proportion of those reads that originated with each animal (sum = 1). For each night, we then summed this value across all 16 resource zones for each animal. For males, this is a measure of |
|  |  | competitive ability. For females, interpretation is difficult, but it is most closely a measure of isolation from other females. Range = 0-2.7, mean = 0.3.<br>See methods for additional details. |
| **Social**<br>**Network**<br>**Phenotypes** | **Edge Strength** | The average strength of the connection between each animal and all other animals that it was connected to in the network each night. A measure of the strength of its average social relationship. Range = 0-4.1, mean = 0.5. |
|  | **Opposite Sex Edge Strength** | The average strength of the connection between each animal and all other animals of the opposite sex that it was connected to in the network each night. A measure of the strength of its average social relationship. Range = 0-3.3, mean = 0.2. |
|  | **Eigenvector Centrality** | A measure of how connected an animal was to other members of the network, taking into consideration the strength of connectivity of each animal that the focal animal was connected to. Range = 0-1, mean = 0.28. |
|  | **Betweenness Centrality** | A measure of centrality that is based on the extent to which an animal lay along the shortest path between other pairs of animals in the network. Range = 0-576, mean = 29. |
|  | **Closeness Centrality** | A measure of centrality based on the length of the average path between any given animal and other animals in the network. Calculated as the inverse sum of the path length (number of connective steps) between the focal animal and each other animal in the network. Range = 0.27-1, mean = 0.61. |

**Table S2.**
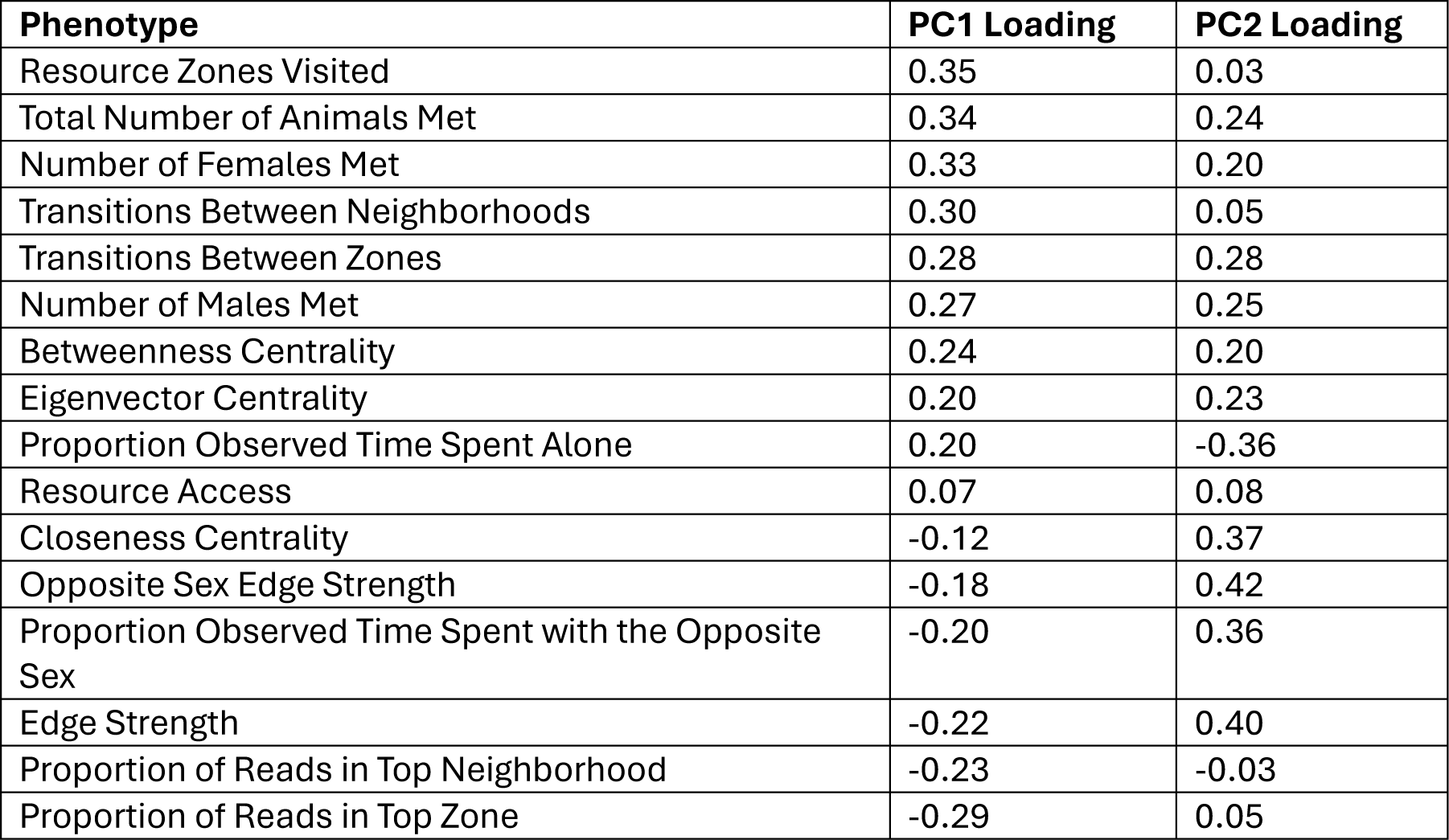
Loadings of each of the 16 phenotypes included in the principal component analysis onto principal components 1 and 2.

**Table S3.** The relationship between average adult resource access and each other adult phenotype measured for each sex.

| Phenotype | Sex | R<sup>2</sup> | p value |
| --- | --- | --- | --- |
| Average Opposite Sex Edge Strength | Male | 0.69 | < **0.0001** |
|  | Female | 0.02 | 0.32 |
| Nightly Females Met | Male | 0.68 | < **0.0001** |
|  | Female | 0.06 | 0.09 |
| Number of Nightly Transitions Between Zones | Male | 0.65 | < **0.0001** |
|  | Female | 0.05 | 0.11 |
| Total Nightly Animals Met | Male | 0.35 | < **0.0001** |
|  | Female | 0.04 | 0.16 |
| Time of First Nightly Transition Between Zones | Male | 0.37 | < **0.0001** |
|  | Female | 0.07 | 0.07 |
| Betweenness Centrality | Male | 0.27 | < **0.0001** |
|  | Female | 0.00 | 0.75 |
| Closeness Centrality | Male | 0.28 | < **0.0001** |
|  | Female | 0.03 | 0.25 |
| Number of Nightly Transitions Between Neighborhoods<sup>^</sup> | Male | 0.22 | **0.002** |
|  | Female | 0.10 | **0.03** |
| Proportion of Nightly Reads in Top Neighborhood | Male | 0.34 | < **0.0001** |
|  | Female | 0.23 | **0.0008** |
| Average Edge Strength | Male | 0.32 | < **0.0001** |
|  | Female | 0.27 | **0.0002** |
| Eigen Vector Centrality | Male | 0.15 | **0.01** |
|  | Female | 0.09 | **0.04** |
| Proportion of Observed Time Spent With a Member of the Opposite Sex | Male | 0.08 | 0.06 |
|  | Female | 0.04 | 0.20 |
| Nightly Males Met | Male | 0.01 | 0.50 |
|  | Female | 0.00 | 0.87 |
| Nightly Resource Zones Visited | Male | 0.00 | 0.78 |
|  | Female | 0.08 | 0.06 |
| Proportion of Nightly Reads in Top Resource Zone | Male | 0.05 | 0.13 |
|  | Female | 0.11 | **0.02** |
| Proportion of Observed Time Spent Alone | Male | 0.16 | **0.009** |
|  | Female | 0.31 | < **0.0001** |

<sup>^</sup>Analysis excludes animal 3059-1, which was an extreme outlier on this metric

**Figure S1.**
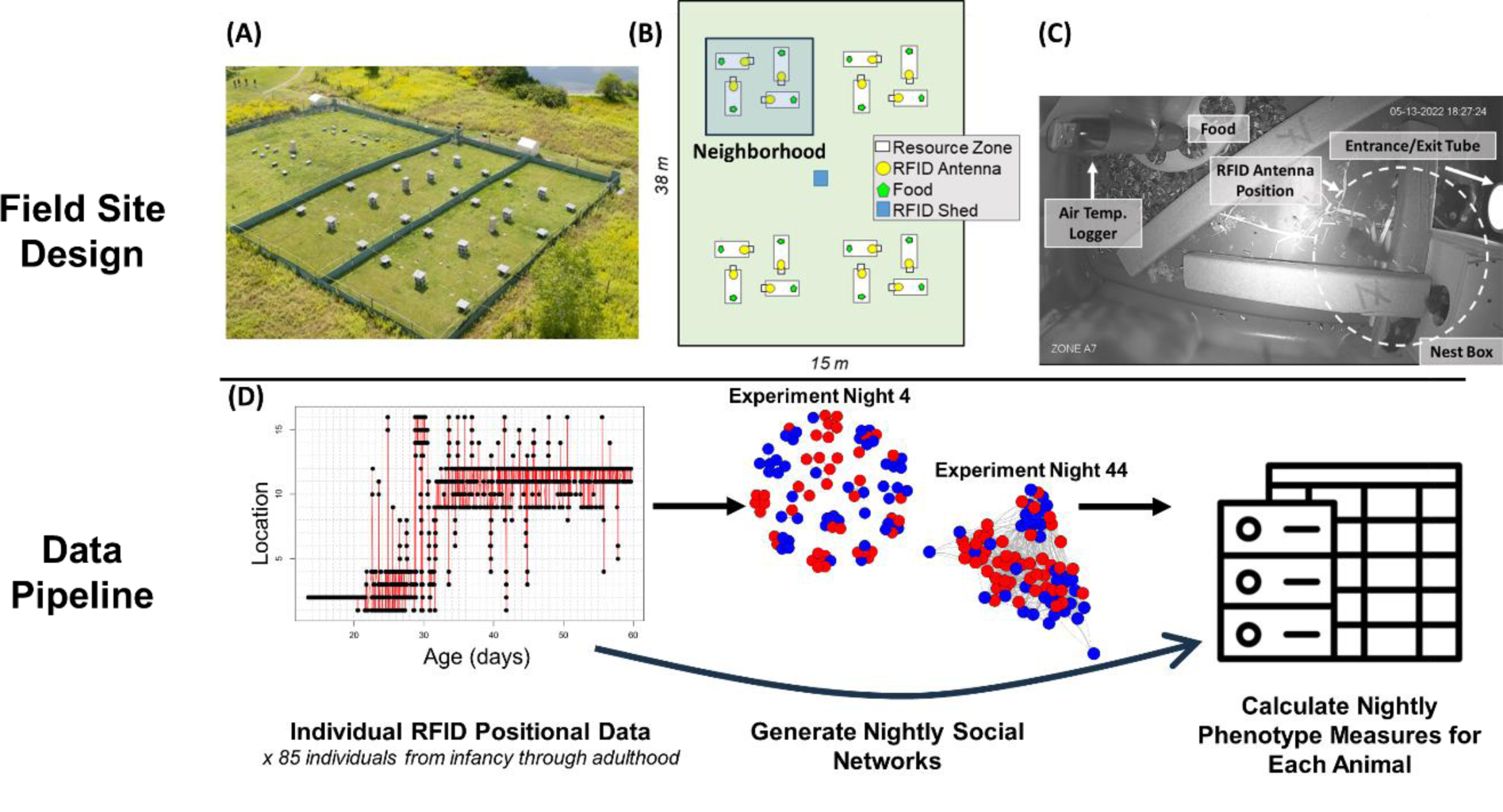
The experimental approach. (A) An aerial view of our field site. Note that the configuration of resource zones in this photograph is different from the configuration in this experiment. (B) Schematic layout of our field enclosure in this experiment (not to scale). We placed litters, along with their mothers inside of the nest box in one of 16 resource zones, which were distributed into four “neighborhoods” of four zones each. (C) An interior view of one of our resource zones. (D) Overview of data processing pipeline for our experiment. Example RFID positional data show the known location of a single individual in our enclosure over the course of the experiment. Each point represents an RFID read at a given resource zone (y-axis). Red lines between points indicate transitions between resource zones.

**Figure S2.**
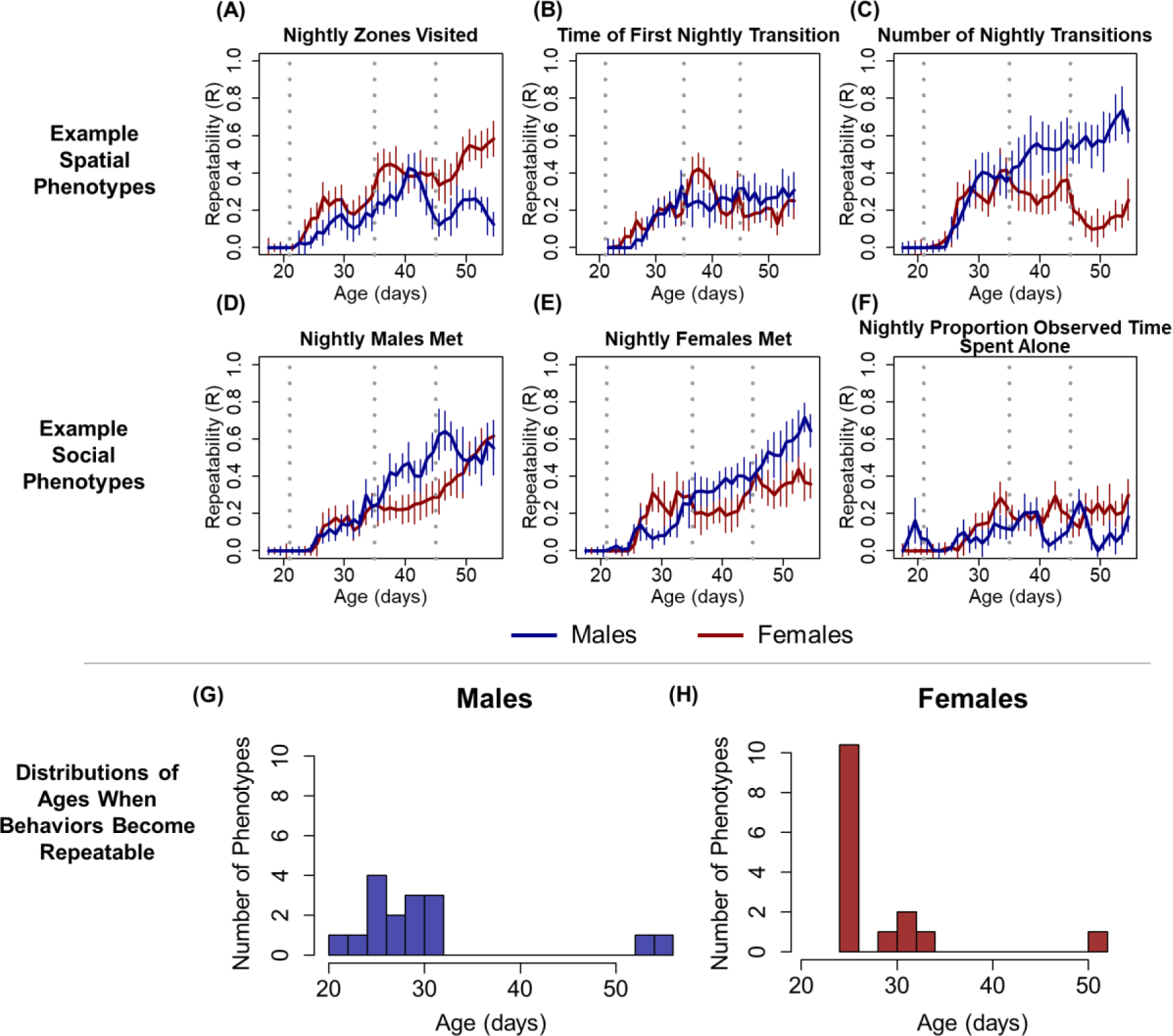
Both males and females developed repeatable individual differences in spatial and social phenotypes, beginning in the juvenile period. (A-F) Examples of repeatability data for four representative spatial and social phenotypes. The repeatability measure on the y-axis controls for maternal/litter identity. Each point represents individual repeatability estimates from each sex during a sliding five-day window, with the x-axis value representing the center of that window. Error bars indicate standard error of repeatability estimates. Vertical dashed lines indicate approximate ages of weaning (21 days), sexual maturity (35 days), and first successful mating (46 days). (G-H) The distributions of the ages at which behaviors became repeatable for males and females.

**Figure S3.**
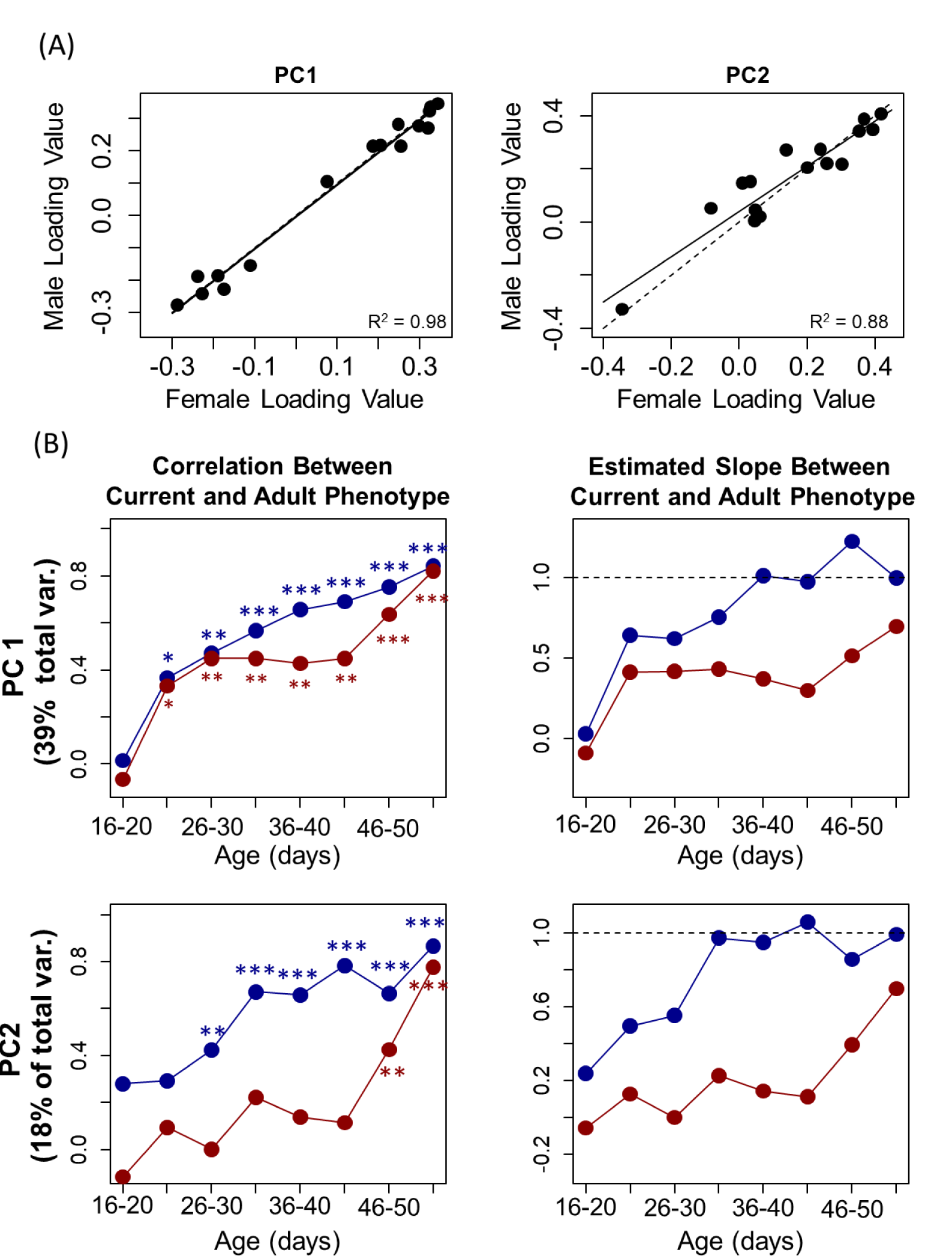
Results in Figure 1 are unchanged if we use a sex-specific principal components approach. (A) Loading coefficients of individual phenotypes are extremely similar in male-specific and female-specific principal components 1 and 2. Solid line indicates modeled correlation, dashed line indicates a one-to-one theoretical ideal. (B) The same analyses as in Figure 1B, except using sex-specific principal components 1 and 2. The correlation between earlier and adult behavior is stronger in males for both PC1 and PC2 and the slope of the relationship between earlier and adult behavior is closer to 1. Asterisks denote significance of the correlations depicted in each point (* p < 0.05, ** p < 0.01, *** p < 0.001).

**Figure S4.**
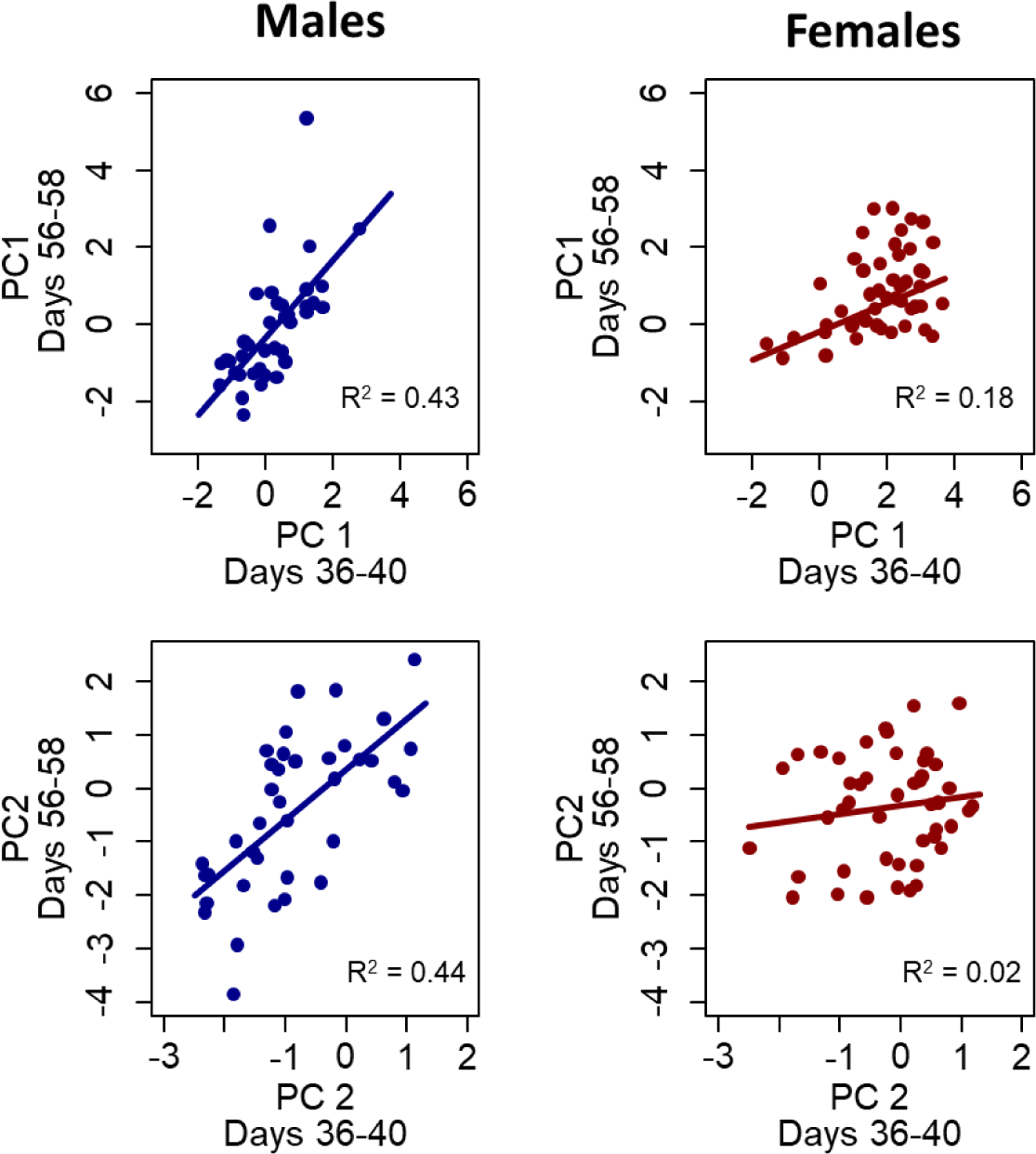
An example of the sets of models contained in Figure 1B. Here we compare the relationship between behavior immediately after sexual maturity (age 36-40 days) and later behavior in adulthood at the end of the experiment (days 56-68 days). For both PC1 (top row) and PC2 (bottom row) the relationship is stronger and the slope estimates are closer to 1.00 for males as compared to females.

**Figure S5.**
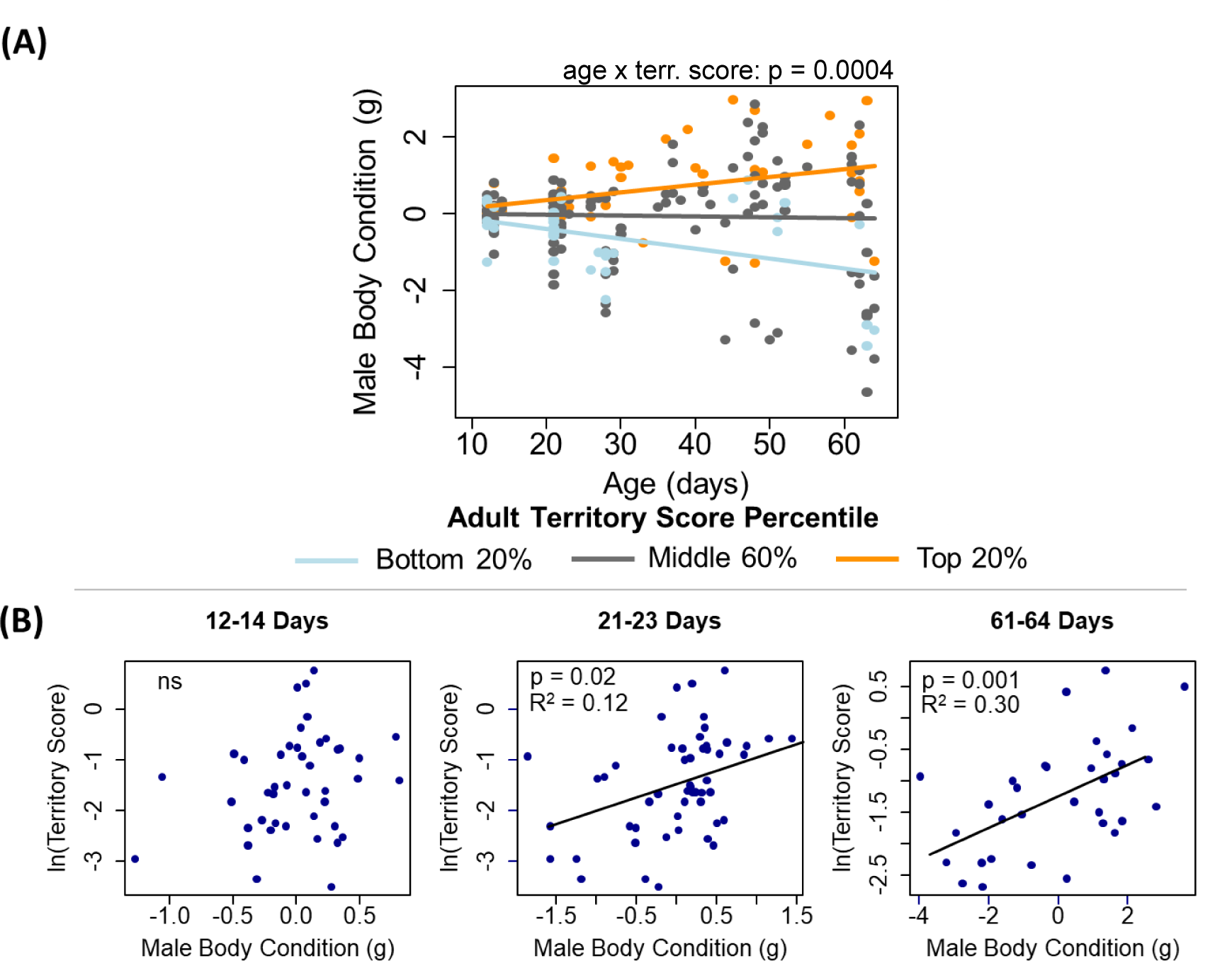
Small differences in initial male body mass are magnified by differential territorial access. (A) The same data as in Figure 3C, but here presented as individual data points. Male adult territory scores are predicted by small differences in body mass in early life, a difference that is magnified over time. The y-axes represent deviations from age-predicted body mass. (B) The strength of the relationship between body condition and adult territory score (days 46-58) increases as males age. Although we collected opportunistic body masses from individuals throughout the experiment, we collected body masses from all animals in the enclosure at three different points: (1) prior to release (age 12-14 days), at weaning (21-23 days), and after we ended the experiment (61-64 days). Initial body condition in infancy did not predict final territory scores (consistent with individuals starting out on an approximately even playing field. However, as males aged, the correlation between territorial behavior and body mass increased in strength, consistent with a competitively induced feedback loop.

